# Synaptic and transcriptionally downregulated genes are associated with cortical thickness differences in autism

**DOI:** 10.1101/208223

**Authors:** Rafael Romero-Garcia, Varun Warrier, Edward T. Bullmore, Simon Baron-Cohen, Richard A.I. Bethlehem

## Abstract

Differences in cortical morphology - in particular, cortical volume, thickness and surface area - have been reported in individuals with autism. However, it is unclear what aspects of genetic and transcriptomic variation are associated with these differences. Here we investigate the genetic correlates of global cortical thickness differences (ΔCT) in children with autism. We used Partial Least Squares Regression (PLSR) on structural MRI data from 548 children (166 with autism, 295 neurotypical children and 87 children with ADHD) and cortical gene expression data from the Allen Institute for Brain Science to identify genetic correlates of ΔCT in autism.

We identify that these genes are enriched for synaptic transmission pathways and explain significant variation in ΔCT. These genes are also significantly enriched for genes dysregulated in the autism post-mortem cortex (Odd Ratio (OR) = 1.11, P_corrected_ < 10^−14^), driven entirely by downregulated genes (OR = 1.87, P_corrected_ < 10^−15^). We validated the enrichment for downregulated genes in two independent datasets: Validation 1 (OR = 1.44, P_corrected_ = 0.004) and Validation 2 (OR = 1.30; P_corrected_ = 0.001). We conclude that transcriptionally downregulated genes implicated in autism are robustly associated with global changes in cortical thickness variability in children with autism.

## Introduction

Autism Spectrum Conditions (henceforth ‘autism’) are characterized by difficulties in social communication alongside unusually narrow interests and restrictive, repetitive behaviours, a resistance to unexpected change, and sensory hypersensitivity ^1^. In addition to behavioural and clinical differences, differences in cortical morphology between individuals with autism compared to typical controls have been reported ^2–5^. While heterogeneous, recent studies have reported increased cortical volumes in the first years of life with autism compared to controls, with accelerated decline or arrest in growth in adolescents ^3,4^. Changes in cortical volume may be attributed to changes in cortical thickness (CT), changes in surface area, or both ^3^. In support of this, studies have separately identified differences in both surface area ^6^ and cortical thickness (CT) ^7^ in children with autism. For example, Smith and colleagues show that the developmentally accelerated gain in grey matter volume in autism is largely driven by the lack of typical age-related CT decrease. Furthermore, earlier studies identified differential trajectories in CT development in autism ^8^ as well as CT difference in autism in specific brain regions ^9,10^. Despite significant heterogeneity in cortical morphology across autism imaging studies ^11^, recent studies have indicated alterations in areas associated with higher cognition (e.g., language, social perception and self-referential processing) ^7,12^. This has been supported by observed differences in cortical minicolumns in association areas in individuals with autism ^13^.

It is unclear what contributes to these differences in cortical morphology in individuals with autism. Genetic factors play a major role in the development of brain networks and volumes in typically developing individuals ^14–16^. For instance, twin heritability of cortical thickness measures suggest modest to high heritability for most regions of the brain ^17^. In parallel, the contribution of genetic factors for autism has been estimated between 50–90% ^18–20^. Different classes of genetic variation have been associated with risk for autism. Several recent studies have identified a significant contribution of rare, *de novo* putative loss-of-function mutations for autism ^21–25^. In addition, common genetic variants, cumulatively, account for approximately half of the total variance in risk for autism ^18^. Studies have also identified genes dysregulated in the autism post-mortem cortex ^26–28^, enriched in processes such as synaptic transmission and astrocyte and microglial genes. These dysregulated genes may either represent causal mechanisms for risk or compensatory mechanisms as a result of upstream biological and cellular changes. Genes dysregulated in the autism postmortem cortex are also enriched in specific gene co-expression modules identified using both adult ^26–28^ and fetal ^29^ cortical post-mortem samples.

Despite considerable progress in understanding neuroanatomical and genetic risk for autism, several questions remain. Mechanistically, it is likely that genetic risk variants alter neuroanatomical structural and functional properties, contributing to behavioural and clinical phenotypes. Given the heterogeneity in autism imaging findings ^11^, it is pertinent to ask how genetic risk for autism is associated with variability in cortical morphology observed in individuals with autism. Thus, the goal of the present study was to identify molecular correlates of disease-related neuroanatomy irrespective of regional specific neuroanatomical differences that may not replicate well across studies ^11^. Here, focusing on CT, we ask 3 specific questions: Q1) Which genes and biological pathways are associated with in cortical thickness variability (ΔCT) in children with autism? Q2) What is the spatial expression profile of genes associated with ΔCT? and Q3) Are these genes enriched for three different classes of risk factors associated with autism: rare, *de novo* variants, common genetic variants, and/or dysregulated genes in the post-mortem cortex? We address these questions by combining analysis of ΔCT in autism, as measured with MRI, with gene expression postmortem data provided by the Allen Institute for Brain Science (AIBS; ^30,31^).

## Methods

### Overview

We first assessed differences in CT (ΔCT) across 308 cortical regions in individuals with autism by extracting cortical thickness estimates for 62 children with autism (cases) and 87 matched typically developing individuals (controls) from the ABIDE-I (Table 1; Discovery dataset). Using median gene expression of 20,737 genes from 6 post-mortem cortical brain samples ^30^, we conduct a Partial Least Squares regression (PLSR), a data reduction and regression technique, to identify significant genes and enriched pathways that contribute to ΔCT (Q1). We next quantified the expression of the same significant genes in terms of their spatial profile by comparing them across the different brain regions and Von Economo classes (Von Economo and Koskinas, 2008), which provides a way of assessing the hypothesis that there would be a global differentiation between higher order cognitive processing and more primary sensory processing (Q2). We tested any significant genes for enrichment for classes of genetic and transcriptomic variation associated with autism (Q3): 1. Genes dysregulated in the autism post-mortem cortex; 2. Adult cortical gene co-expression modules associated with dysregulated genes in the autism post-mortem cortex; 3. Fetal cortical gene co-expression modules associated with dysregulated genes in the autism postmortem cortex; 4. Genes enriched for rare, de novo loss of function mutations in autism; and 5. Common genetic variants associated with autism. To assess the replicability of the findings, we validate the results using two independent datasets from ABIDE-II (Table 1). In parallel, we also used a second list of genes dysregulated in autism identified using a partially-overlapping cortical gene expression dataset of autism and control post-mortem brain samples to validate the enrichment analysis across all datasets. To assess specificity of our results, we furthermore sought to answer these questions in a matched MRI dataset of children with ADHD, another childhood psychiatric condition. A schematic overview of the study protocol is provided in Figure 1.

**Figure 1:**
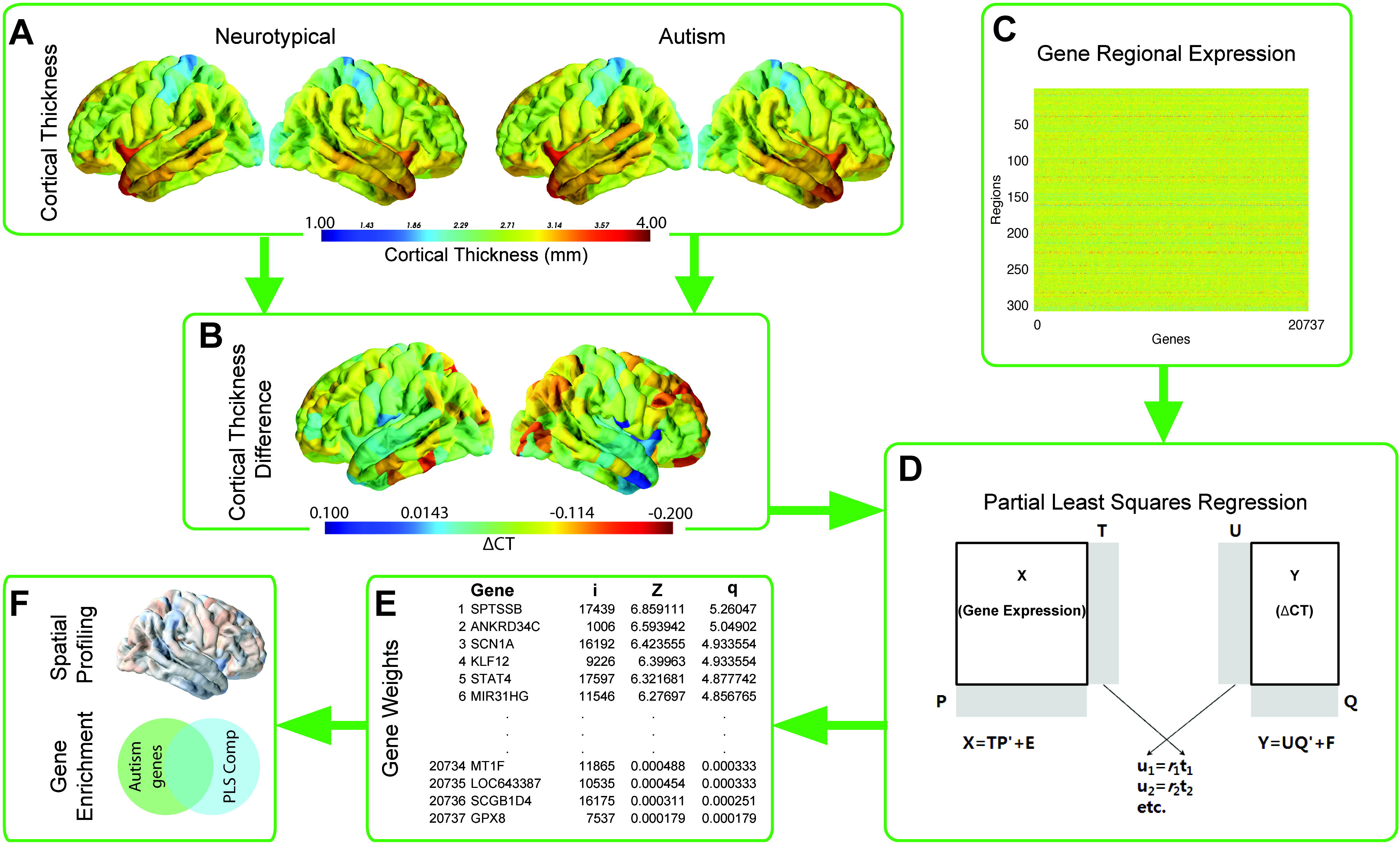
Schematic overview of the methodology used to identify gene contribution. Mean cortical thickness was extracted for both the autism and the neurotypical groups across 308 cortical nodes (Panel A). A difference score in cortical thickness (ΔCT; autism - neurotypical) was calculated between these two groups (Panel B). In parallel the median AIBS gene expression profiles for 20,737 genes were calculated across the same 308 cortical nodes used in the imaging analysis (Panel C). Both these streams were included in a bootstrapped PLSR analysis that used the gene expression profiles as predictors and the ΔCT as response variable (Panel D). The PLSR assigns weights to each gene in terms of its contribution to the overall model in each component. Bootstrapped standard errors were derived and the gene weights were Z-transformed and corrected for multiple comparison using a Benjamini and Hochberg FDR correction (Panel E; i = gene index number, z = z-score for that gene’s association and q = FDR corrected z-score). Genes that were significant after FDR correction (z-score>1.96) were analysed in terms of their spatial expression as well as tested for enrichment against three classes of risk for autism: dysregulated autism genes in the postmortem cortex, genes harbouring rare de-novo variants, and common genetic variants in autism (Panel F).

**Table 1.** Table 1: Descriptive statistics for all four datasets. The Discovery cohort was obtained from ABIDE-I. The validation cohorts were obtained from the ABIDE-II (Validation 1: Georgetown University, Validation 2: Kennedy Kreiger Institute). The n-row denotes the number of subjects with the number of females (F) provided in parenthesis, FIQ denotes the full-scale IQ, with standard deviations in parenthesis below. *Indicates that the same controls were used for both the Autism Discovery and the ADHD datasets. Further details on the Discovery and ADHD datasets are described elsewhere ^32^.

|  | Discovery |  | Validation 1 |  | Validation 2 |  | ADHD |  |
| --- | --- | --- | --- | --- | --- | --- | --- | --- |
|  | Autism | Controls* | Autism | Controls | Autism | Controls | ADHD | Controls* |
| **n** | 62 | 87 | 48 | 54 | 56 | 154 | 69 | 87 |
|  | (0 F) | (0 F) | (8 F) | (27 F) | (15 F) | (56 F) | (0 F) | (0 F) |
| **Age** | 10.07 | 10.04 | 10.98 | 10.43 | 10.32 | 10.34 | 9.99 | 10.04 |
|  | (±1.11) | (±1.13) | (±1.53) | (±1.71) | (±1.51) | (±1.20) | (±1.17) | (±1.13) |
| **FIQ** | 108.86 | 110.98 | 118.68 | 122.04 | 103.42 | 114.4 | 107.95 | 110.98 |
|  | (±16.94) | (±10.39) | (±15.01) | (±13.27) | (±15.99) | (±10.55) | (±14.18) | (±10.39) |

### Discovery dataset

#### Neuroimaging, gene expression and PLS regression

Discovery imaging data used in this study are described in detail in the supplementary materials and elsewhere ^32^. In short, structural T1 weighted MPRAGE images were obtained from the ABIDE I database (http://fcon_1000.projects.nitrc.org/indi/abide/, selecting participants in the age range from 9-11, all males. All subjects were matched on age, and IQ between groups (see Table 1; Discovery Data) (see ^32^ and Supplementary Materials for details on matching and scanner site). CT estimates were extracted using freesurfer and visually inspected for quality of segmentation by two independent researchers. Only when there was consensus between researcher were images included. Next, images were parcellated into 308 cortical regions and mean CT for these regions was extracted. In addition, scanner site was regressed out from CT estimates and the residuals were added to the group mean to allow for easier interpretation. The final sample consisted of 62 children with autism (cases) and 87 neurotypical individuals (controls).

We used the transcriptomic dataset of the adult human brain created by the Allen Institute for Brain Science (http://human.brain-map.org) ^30,31^. The anatomical structure associated to each tissue sample was determined using the MRI data provided by the AIBS for each donor. The high-resolution parcellation with 308 cortical regions, employed in the neuroimaging dataset, was warped from the anatomical space of the average subject provided by FreeSurfer (fsaverage) into the surface reconstruction of each AIBS donor brain. After pre-processing regional gene expression values were represented as a 308×20737 matrix that contained the whole-genome expression data for the 308 MRI regions of interest. Code used to determine regional gene expression levels is available at (https://github.com/RafaelRomeroGarcia/geneExpression_Repository) and data used can be downloaded from Cambridge Data Repository ^33^. More details on tissue sample handling, processing, batch correction and consistency of gene expression data across donors are provided in the Supplementary Materials.

We used partial least squares regression (PLSR) to identify which genes were significantly associated with ΔCT. After obtaining PLS weights for each gene, these were z-transformed (based on standard errors obtained from bootstrapping) and FDR-adjusted using a Benjamini-Hochberg FDR correction with alpha set at 0.05. Only genes that passed FDR correction were included in enrichment analysis. We used significant genes with both negative and positive weights in our analysis. As our dependent variable, ΔCT, had both positive and negative values, weight signs were not informative about directionality in the analysis (in short: a more positive association would mean a bigger ΔCT in either direction, similarly a more negative weight would also imply a bigger ΔCT in either direction). A detailed description of the PLSR regression and the detailed rationale behind choosing the unsigned weights is provided in the Supplementary Materials.

### Genetic modules and enrichment analyses

We used Enrichr (http://amp.pharm.mssm.edu/Enrichr) ^34,35^ for enrichment of significant PLSR genes for each component against Gene Ontology Biological Processes and report significant results after Benjamini-Hochberg FDR correction (q < 0.05). Cell-type specific enrichment was conducted for five broad classes of cells: neurons, astrocytes, oligodendrocytes, microglia, and vascular cells ^36^. We defined cell-type specific genes as the top 500 genes with higher expression in the cell-type compared to the remaining five genes.

As these classes of genes are distinct with minimal overlap, we used a Bonferroni correction to correct for cell-type specific enrichment.

We also investigated the enrichment in different classes of risk genes using logistic regression (more detail on each class of genes can be found in the supplementary materials):

1. *Transcriptionally dysregulated genes* (n = 1143, 584 upregulated and 558 downregulated in the autism cortex) were identified from Parikshak *et al.* (2016).
2. *Adult gene co-expression modules* (Parikshak et al., 2016).
3. *Fetal gene co-expression modules* (Parikshak et al., 2013).
4. *Genes harbouring rare, de novo, putative loss of function variants* (rare, de novo genes, n = 65) were identified from Sanders et al., (2015).
5. *Common genetic variants* associated with autism were downloaded from the latest data freeze from the Psychiatric Genomics Consortium (5,305 cases and 5,305 pseudocontrols). Gene based P-values and Z scores were obtained using MAGMA for each gene ^37^.

Enrichment analyses for the different classes of autism risk genes were done using logistic regression after accounting for gene length as a covariate. Enrichments are reported as significant if they had a Benjamini-Hochberg FDR adjusted P-value < 0.05 ^38^ and if they have an enrichment odds ratio (OR) > 1. The supplementary material provides further details about the gene sets and the methods used.

### Validation and specificity

We conducted extensive validation of our initial results against two independent datasets and checked for specificity of an autism effect against a matched ADHD dataset. There are significant phenotypic and genetic correlations between the two conditions, and we had access to MRI data from children with ADHD ^32^, making this a suitable dataset for testing specificity. Details on all these three datasets are provided in the supplementary materials.

## Results

### Autism discovery MRI dataset

#### PLSR analyses and characterization

Cross-validation using an initial 35 component analyses identified that a 13-component model had the best fit (Supplementary Table S1). Note that the number of components chosen for the model does not affect the individual component composition. Consequently, PLSR was run using a 13-component model. Four components (Components 1, 3, 4 and 6) explained more than 10% of the variance (Supplementary Figure S1). However, variance in ΔCT explained by PLS components was higher than expected by chance *only* for the first component (P = 0.009, 10,000 permutations) but not for the remaining components (P = 0.303, P = 0.693 and P = 0.394, for components 3, 4 and 6, respectively). Thus, *only* PLSR1 was used for subsequent analyses and we only included genes that passed FDR correction (q < 0.05). Only the GO term “Synaptic Transmission” in component 1 (PLSR1) survived FDR correction for multiple comparisons (P_corrected_ = 0.00006). PLSR1 was also significantly enriched for 11 pathways (Table S2) in the Kyoto Encyclopedia of Genes and Genomes (KEGG). There was a significant positive correlation between ΔCT and the scores of PLSR1 (r = 0.32; P = 4.15×10^−9^).

Cell-type specific analysis identified a significant enrichment for neurons (OR = 1.1; P_corrected_ = 3.19×10^−12^), but no enrichment for genes enriched in astrocytes (OR = 1; P_corrected_ = 1), oligodendrocytes (OR = 0.99; P_corrected_ = 1), microglia (OR = 0.97; P_corrected_ = 0.43) or vascular cells (OR = 0.97; P_corrected_ = 0.62).

#### Topographical enrichment analyses

Previous studies reported an association between cortical thickness and cytoarchitectural cortical features ^39^ linked to specific abnormalities in laminar thickness of supragranular layers-of the cortex of schizophrenia patients ^40^. Here, we also conducted spatial characterization of PLSR1 across all 5 Von Economo classes ^41^ as well as an additional 2 subtype classes covering limbic regions and allocortex (class 6) and insular cortex (class 7) ^15,42^. We expected a potential differentiation between higher order cognitive processing and more primary sensory processing. The genes in PLSR1 were significantly over-expressed in secondary sensory and association cortices (VE classes 2, 3 and 4: all P_corrected_ < 0.01) compared to a null distribution. In limbic and insular regions, however these genes appeared to be under-expressed (VE classes 6 and 7: all P_corrected_ < 0.01). However, they also appear to be over-expressed in granular and primary motor cortices (VE Class 1). Figure 2 shows the results from the spatial characterization of the first component across all VE classes.

**Figure 2:**
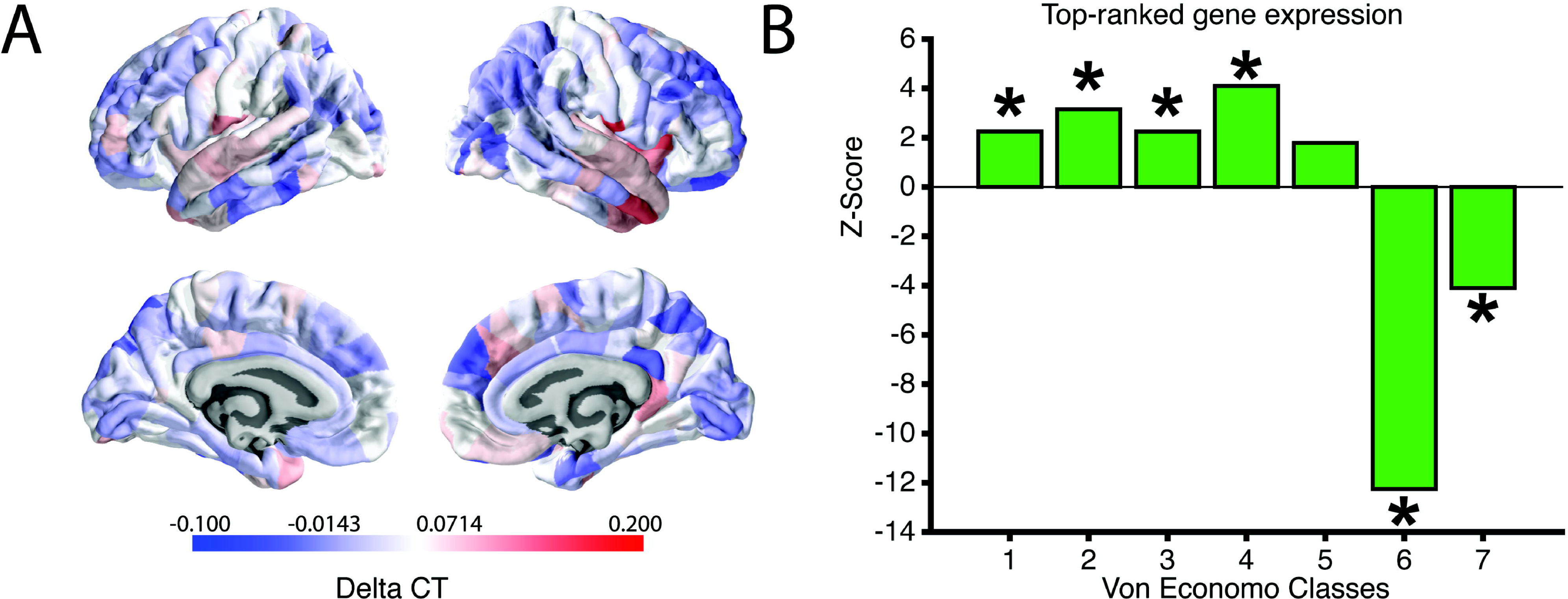
Expression and Von Economo classification for PLSR1. The heatmap in panel A shows the ΔCT distribution across all 308 cortical regions. The barplot in panel B shows the z-scores of the mean distribution across the different Von Economo Classes (Class 1: granular cortex, primary motor cortex. Class 2: association cortex. Class 3: association cortex. Class 4: dysgranular cortex,secondary sensory cortex. Class 5: agranular cortex, primary sensory cortex. Class 6: limbic regions, allocortex. Class 7: insular cortex.). All significant over-or under-expression classes are marked with an asterisk. To determine significance, we used permutation testing and an False Discovery Rate corrected p-value < 0.025 to fully account for two-tailed testing.

#### Gene enrichment analyses

We identified a significant enrichment for genes that are dysregulated in the autism post-mortem cortex (OR = 1.21; P_corrected_ < 2.81×10^−15^), driven entirely by genes downregulated in autism cortex (OR = 1.87; P_corrected_ < 3.55×10^−16^). In comparison, there was no enrichment for upregulated genes (OR = 1.01; P_corrected_ = 0.49). The downregulated genes have been previously reported to be enriched for several GO terms including synaptic transmission ^26^.

Transcriptionally dysregulated genes can reflect several different underlying processes. To provide better resolution of the processes involved, we next investigated if this enrichment was associated with six adult co-expression modules associated with dysregulated autism genes ^26^. Three of these were associated with genes downregulated in the autism postmortem cortex (M4, M10, M16), and three were enriched for genes upregulated in the autism post-mortem cortex (M9, M19, and M20) compared to controls. As we had identified a significant enrichment for downregulated autism genes but not for the upregulated autism genes, we hypothesized that gene co-expression modules associated with downregulated genes would also be enriched for association with PLSR1 genes. Indeed, PLSR1 was enriched for all three downregulated modules but none of the 3 upregulated modules. See Figure 3J, Table 2 and supplementary Table S6.

We also investigated if the significant genes in PLSR1 were enriched in specific cortical developmental modules ^29^. The Mdev13, Mdev16, and the Mdev17 modules are enriched for transcriptionally dysregulated genes in autism postmortem frontal and temporal cortices ^29^. The Mdev2 and the Mdev3 modules are enriched for rare variants identified in autism ^29^. Again, we identified significant enrichment for 3 adult co-expression modules enriched for transcriptionally dysregulated genes. For the two modules associated with rare, *de novo* variants, we identified fewer PLSR1 genes than expected by chance. See Figure 3J, Table 2 and Supplementary Table S6. We did not identify a significant enrichment for rare, *de novo* genes. We also did not identify a significant enrichment for common variants using MAGMA to collapse SNP based P-values to gene based P-values (OR = 1.00; P_corrected_ = 0.29). Results of the gene enrichment analysis are provided in Figure 3J, Table 2 and Supplementary Table S6.

**Table 2:** Gene enrichment. Odds ratio scores, confidence intervals and significance of all major classes of gene enrichment investigated in the discovery dataset.

| Category | Dataset | OR | Upper<br>CI<br>(95%) | Lower<br>CI<br>(95%) | P | P<sub>corrected</sub> |
| --- | --- | --- | --- | --- | --- | --- |
| Autism transcription | Dysregulated | 1.21 | 1.23 | 1.19 | 1.76E-15 | 2.81E-15 |
| Autism transcription | Downregulated | 1.87 | 1.94 | 1.8 | 2.00E-16 | 3.55E-16 |
| Autism transcription | Upregulated | 1.01 | 1.02 | 1 | 4.99E-01 | 4.99E-01 |
| Adult co-expression modules | Mod4 | 1.08 | 1.08 | 1.07 | 2.00E-16 | 3.55E-16 |
| Adult co-expression modules | Mod10 | 1.07 | 1.08 | 1.07 | 2.00E-16 | 3.55E-16 |
| Adult co-expression modules | Mod16 | 1.08 | 1.08 | 1.07 | 2.00E-16 | 3.55E-16 |
| Adult co-expression modules | Mod9 | 0.93 | 0.94 | 0.92 | 2.01E-14 | 2.92E-14 |
| Adult co-expression modules | Mod19 | 0.93 | 0.94 | 0.92 | 2.00E-16 | 3.55E-16 |
| Adult co-expression modules | Mod20 | 0.97 | 0.97 | 0.96 | 6.22E-05 | 7.66E-05 |
| Common variants | Common<br>variants | 1 | 1.01 | 1 | 2.75E-01 | 2.93E-01 |
| Rare variants | Rare variants | 0.96 | 0.99 | 0.93 | 2.42E-01 | 2.76E-01 |
| Fetal co-expression modules | Moddev2 | 0.97 | 0.97 | 0.96 | 1.28E-11 | 1.70E-11 |
| Fetal co-expression modules | Moddev3 | 0.96 | 0.97 | 0.96 | 2.00E-16 | 3.55E-16 |
| Fetal co-expression modules | Moddev13 | 1.04 | 1.04 | 1.04 | 2.00E-16 | 3.55E-16 |
| Fetal co-expression modules | Moddev16 | 1.06 | 1.06 | 1.05 | 2.00E-16 | 3.55E-16 |
| Fetal co-expression modules | Moddev17 | 1.04 | 1.05 | 1.04 | 2.00E-16 | 3.55E-16 |

**Figure 3:**
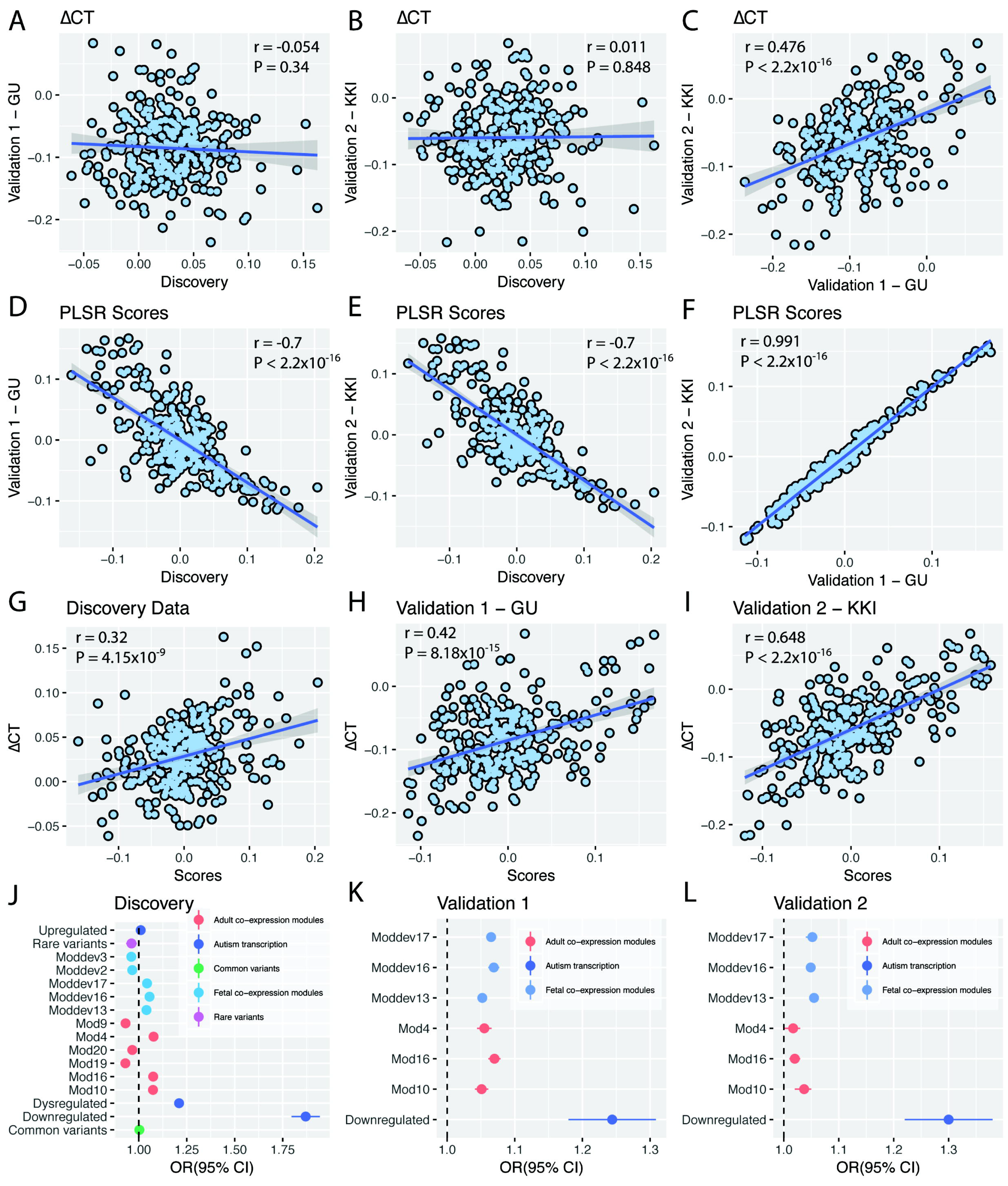
Gene enrichment and dataset comparisons. Panels A-C show the correlation between ΔCT in the three datasets. Panels D-F show the correlation between the PLSR scores of all three datasets. Panels G-I show the correlation between ΔCT and the PLSR scores in all three datasets (indicating that increased scores are strongly correlated with increased ΔCT). Panel J shows the odds ratios for the gene-enrichment analysis in the discovery dataset. All significantly enriched modules were replicated in the validation datasets (panels K and L) apart from module 4 of the adult co-expression modules. Pearson correlation coefficient and P-values of the correlations are provided in the top of the respective panels.

### Validation of initial findings

#### PLSR analyses and characterization

We validated all analyses using ΔCT from two independent cohorts (Table 1). There was a non-significant correlation in ΔCT between the discovery and the two validation datasets (Figure 3A and 3B), which is in line with recent large scale assessments of autism neuroimaging studies ^11^. This may be explained by factors such as heterogeneity due to scanner sites in the discovery dataset, age of onset of puberty, and clinical conditions. There was a significant positive correlation in ΔCT between the two validation datasets (r = 0.476; P < 2.2×10^−16^). Heterogeneity in autism neuroimaging studies is well documented and complex ^11,43^, but it should be emphasized that the present analysis focuses on the relation between whole-brain variation in ΔCT and whole-brain variation in gene expression, thus a lack of spatial overlap in ΔCT does not affect the ΔCT – Gene relation.

Again, only the first component (PLSR1-validation1 and PLSR1-validation2) (P < 10^−14^, 10,000 permutations) (Supplementary Figure S2) explained a significant amount of the variance. There was a significant positive correlation between ΔCT and the gene expression scores in both validation datasets (Figures 3H and 3I). Further, PLSR1 was enriched for the GO term ‘Synaptic transmission’ (P_corrected_ = 1×10^−4^ for both Validation 1 and Validation 2). In addition, for Validation 2, the PSLR component was also enriched for the GO term ‘Membrane depolarization’ (P_corrected_ = 7×10^−4^). KEGG pathway enrichment for both datasets are provided in Supplementary Tables S3 and S4. Further, we replicated the cell-type enrichment in genes expressed in neurons (OR = 1.13; P = 2.01×10^−7^ for Validation 1 and OR = 1.06; P = 0.009 for Validation 2).

**Figure 4:**
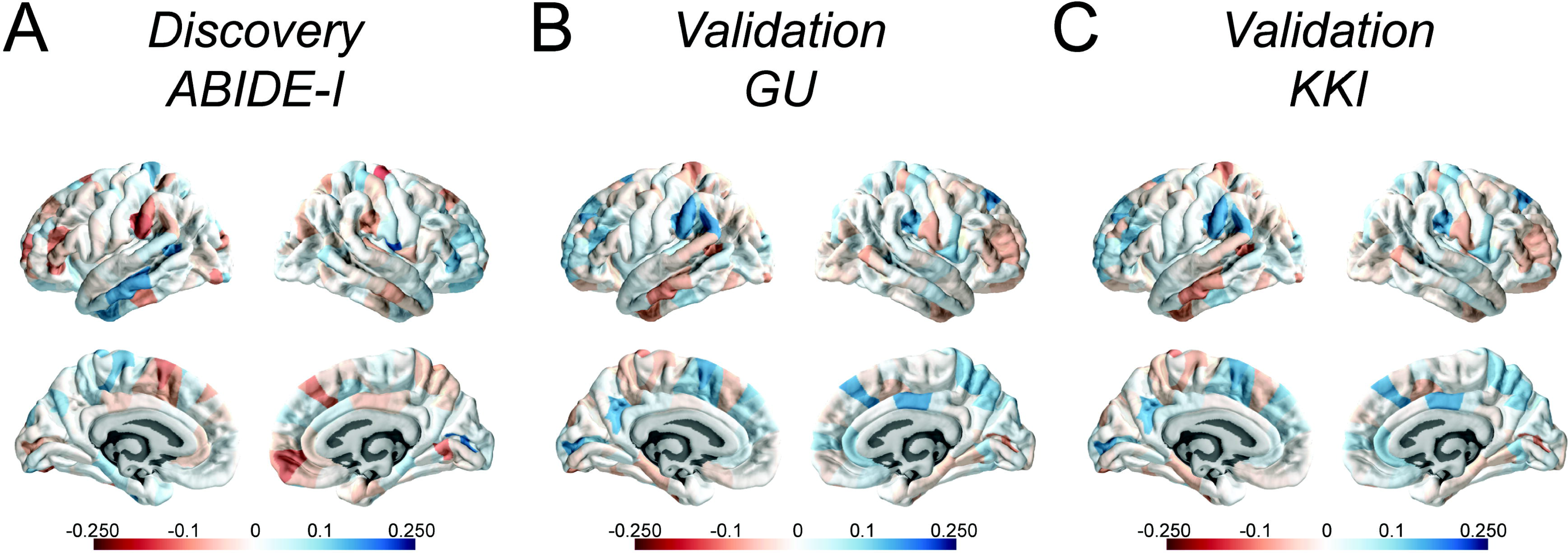
PLSR1 scores for all autism datasets. Panels A to C represent the PLSR1 scores for the three autism datasets across 308 cortical regions. A represents the discovery dataset, B represents Validation 1 and C represents Validation 2

#### Gene enrichment analyses

We replicated the significant enrichment of transcriptionally downregulated genes in the autism post mortem cortex (Validation 1: OR = 1.24; P_corrected_ = 0.004; Validation 2: OR = 1.3; P_corrected_ = 0.001), providing confidence in the robustness of our initial results (Figures 3K and 3L and Supplementary Table S6). Mirroring the enrichment with the downregulated genes in the autism post-mortem cortex, we also identified enrichment for the three adult gene co-expression modules that are enriched for downregulated genes in Validation 1 (M4, M10, and M16) and two of the three (M10 and M16) adult gene co-expression modules for Validation 2 (Figures 3K and 3L, Supplementary Table S6). We also replicated the enrichment for the three fetal gene co-expression modules (Mdev13, Mdev16, and Mdev17) in both validation datasets (Figures 3K and 3L, Supplementary Table S6). Non-significant co-expression modules remained non-significant in the validation datasets.

#### Comparison with ADHD

PLSR analysis of ΔCT in ADHD data did not identify any components that significantly explained the variance in ΔCT (Supplementary Table S5). Thus, we did not consider the ADHD dataset for further analyses. Details of the number of components, the model fit, and the variance explained are provided in the supplementary materials.

## Discussion

Here we report the association of transcriptionally downregulated genes in the autism post-mortem cortex with global differences in cortical thickness in 166 children with autism and 295 neurotypical children. Using partial least squares regression on a discovery dataset of 62 cases and 87 controls, we identify one component (PLSR1) that explains a significant proportion of variance in ΔCT and is enriched for the GO term ‘Synaptic Transmission’ and for neuronal genes. This component was enriched for genes downregulated in the autism post-mortem cortex and validated using two independent datasets. We also find that PLSR1 genes are enriched for fetal and adult developmental cortical modules that have been previously reported to be enriched for transcriptionally dysregulated genes in the postmortem autism cortex and for genes involved in synaptic transmission ^26,29^. We were unable to identify genes associated with ΔCT in ADHD, another childhood condition. Our study provides robust evidence linking disease-related variance in CT to synaptic genes and dysregulated genes in the autism post-mortem cortex, linking molecular and macroscopic pathology.

Validation using two independent autism MRI datasets suggests that the results are valid even using MRI data from different cohorts that had different scanner settings. The results were valid despite non-significant correlations in global ΔCT between the discovery and the two validation datasets and sex did not modulate any of the observed differences between datasets (see Supplementary material). This suggests that the same sets of genes are associated with ΔCT regardless of sex. Studies have identified differences in cortical morphology between neurotypical males and females and between males and females with autism ^2,44^. Here we identified a high correlation between a males-only dataset and two males and females combined MRI datasets for the gene weights and gene scores in the first PLS component.

Changes in CT may be due to a host of factors such as changes in myelination, synaptic pruning, and dendritic arborisation. Evidence from rare genetic variants ^45,46^ and transcriptionally dysregulated genes in autism have highlighted a role for synaptic transmission in the aetiology of autism ^26,27^. Transcriptional dysregulation may reflect either a causative risk mechanism for autism, or a compensatory consequence of genetic, hormonal, and environmental risk for autism. Here, we are unable to disentangle if transcriptionally dysregulated genes causally contribute to cortical morphology changes, or if they are both downstream of a common risk mechanism, or both. It is possible that both cortical thickness variability and transcriptional dysregulation are downstream processes of genetic variation implicated in autism, and, as such the enrichment for transcriptionally dysregulated genes need not be causative of cortical morphological changes.

We did not identify enrichments for rare, *de novo* loss of function genes or common variants implicated in autism. The lack of enrichment with rare, *de novo* loss of function genes may be due to both the relative low frequencies of such variants and small proportion of variance in liability explained by rare de novo variants ^18^. In contrast, the lack of enrichment with common variants may be explained by the lack of statistical power of the largest available autism GWAS dataset. Indeed, there is no enrichment for common genetic variants associated with autism in co-expression modules enriched for transcriptionally dysregulated genes in autism ^26^. In contrast, common variants for schizophrenia are enriched in co-expression modules associated with dysregulated genes in schizophrenia ^47^. It is likely that larger samples will better reveal the role of common genetic variants in cortical morphology differences in autism. While we do not know the genetic make-up of the cases and controls, our results likely represent common downstream convergence of upstream genetic perturbations.

In addition, animal studies have shown that several candidate genes for autism risk are regulated by synaptic activity, leading to the hypothesis that dysregulation in synaptic homeostasis is a major risk for autism ^46^. The effects of this can contribute to both processing of input, and to more morphological changes in neuroanatomy via processes such as activity dependent synaptic pruning and dendritic arborization. Post-mortem studies of the brains of children and adolescents with autism have indicated a lack of synaptic pruning ^48^. Investigating the specific role of synaptic genes in altering neural circuitry and cortical morphology will help elucidate the precise molecular mechanisms underlying cortical thickness differences seen in autism.

There are, some caveats that need to be taken into consideration while interpreting these results. Gene expression data was derived from only six post-mortem adult brain samples. Gene expression is known to vary with age ^49,50^. Unfortunately, we are restricted in using the adult gene expression data from the AIBS for several reasons. First, this is the most spatially detailed dataset of gene expression. Second, the availability of MNI coordinates in the adult gene expression datasets allows for mapping of gene expression in distinct brain regions to cortical thickness differences extracted from MRI scans. Third, gene expression changes with age are limited and restricted to specific brain regions. A recent study identified only 9 genes significantly altered globally across the 10 regions investigated in post-mortem tissue samples ^51^, largely driven by glial genes. Cell specific enrichment in our dataset implicated neuronal genes only. Fourth, as autism is a developmental condition, investigating differences in cortical morphology at an early age is important to limit the role of environmental factors that contribute to differences in cortical morphology later in life ^8,52^. Fifth, enrichment for gene expression modules associated with autism risk in the developing cortex provides further confidence that the genes identified here are relevant across the age-spectrum. We do acknowledge that investigating a paediatric specific gene-expression dataset will help further refine the analyses, once this data becomes available.

Lastly, the present study used cortical thickness in contrast to other morphological features such as cortical volume. It is known that grey matter volume relies on the relationship between two different morphometric parameters, cortical thickness and surface area that are unrelated genetically ^53^ and associated with different developmental trajectories ^54^. The combination of at least two different sources of genetic and maturational influence into a unique descriptor of cortical volume would act as a confounding factor that would complicate meaningful analysis of associated genetic weights.

To our knowledge, this is the first study linking different genetic risk mechanisms in autism with changes in cortical morphology. In sum, we have shown that genes that are enriched for synaptic transmission and downregulated in individuals with autism are significantly associated with global changes in cortical thickness. We also show that these genes are generally overexpressed in association cortices. We could validate the results in multiple independent datasets but not in a matched MRI dataset that included individuals with ADHD, showing both replicability as well as selectivity.

## Acknowledgements

This study was funded by grants from the Medical Research Council, UK, the Templeton World Charity Foundation, the Autism Research Trust, and the Wellcome Trust. We thank Luke Kweku Abraham for help with interpretation of the PLSR gene weights. We thank Dr. Elijah Mak and František Váša for help with quality control of the MRI images. We thank Dr. Neelroop Parikshak for help with regression-based enrichment analyses and Prof. Dan Geschwind for valuable comments on the manuscript. RRG was funded by the NeuroScience in Psychiatry Network, Wellcome Trust. VW was funded by St. John’s College, the Cambridge Commonwealth Trust. RAIB was funded by the Medical Research Council, Autism Research Trust, Pinsent Darwin Trust and Cambridge Trust. SBC is supported by the National Institute for Health Research (NIHR) Collaboration for Leadership in Applied Health Research and Care East of England at Cambridgeshire and Peterborough NHS Foundation Trust. ETB is employed half-time by the University of Cambridge and halftime by GlaxoSmithKline (GSK); he holds stock in GSK. The views expressed are those of the author(s) and not necessarily those of the NHS, the NIHR, GSK or the Department of Health.

